# Immunological response against bovine viral diarrhoea virus (BVDV) types 1 and 2 after vaccination with DIVENCE®, measured by ELISA and serum neutralisation on serum and milk samples

**DOI:** 10.1101/2024.08.06.606756

**Authors:** C. Montbrau, M. Gibert, E. Taberner, M. Tapiolas, R. Teixeira, A. Prenafeta

**Affiliations:** Hipra Scientific S.L.U., Avda. La Selva 135, 17170 Amer, Spain; Hipra, Avda. La Selva 135, 17170 Amer, Spain

**Keywords:** BVDV, DIVA, antibody response, neutralising antibodies, ELISA, bulk tank milk, diagnostic, vaccine

## Abstract

**Background:** A novel subunit vaccine, DIVENCE®, comprises different bovine respiratory and reproductive antigens, including BVDV-1 E2 recombinant glycoprotein and BVDV-2 E2 recombinant glycoprotein. This study evaluated the immune response against BVDV-1 and 2 generated by DIVENCE® over the long term in different strains.

**Methods:** Two different studies were conducted to assess the immune response induced by DIVENCE®. In the first study, 20 seronegative young calves (55-104 days of age) were randomly distributed into the vaccinated or control group. All animals were given four intramuscular doses (D0, D21, D204 and D570) of 2 mL DIVENCE® or phosphate-buffered saline (PBS). Blood samples were collected to assess BVDV serum antibodies (ELISA) and neutralising antibodies (SN) against BVDV-1 and BVDV-2. In the second trial, heifers (from 10 months of age until calving) and cows (between first and seventh calving) were included and assigned to the vaccinated and control groups. Three different farms were enrolled in the study. The administration regimen was the same as described in the first study. Blood samples were collected from 32 random animals on each farm (16 vaccinated and 16 control animals). Additionally, bulk tank milk samples from each farm were obtained at different time-points and individual milk samples were obtained from vaccinated and control animals. The immune response against BVDV was analysed in serum and milk samples using different ELISA kits. Neutralising antibodies induced by DIVENCE® against different BVDV isolates from Europe and the Americas were also assessed in this trial.

**Results:** Overall, DIVENCE® induced high levels of total antibodies (ELISA) and neutralising antibodies against BVDV-1 and BVDV-2. These values were significantly (*p*<0.05) greater than the control group from 21 days after the second dose (D42) until the end of the study (D591). The high antibody levels, particularly after the third dose, were similar to those that described efficacy against experimental challenges of BVDV-1 and 2 in pregnant animals. In the field trial, similar results were observed in terms of total antibodies (ELISA) against BVDV, and no induction of anti-p80 antibodies was observed in vaccinated animals at any time. Additionally, analyses on individual and bulk tank milk samples also confirmed that no anti-p80 antibodies were induced in vaccinated animals. Furthermore, high levels of neutralising antibodies were observed against different BVDV isolates from Europe and the Americas.

**Conclusion:** The DIVENCE® vaccine induced a strong immune response against BVDV-1 and BVDV-2, and this response allows infected and vaccinated animals to be differentiated (DIVA vaccine).

## INTRODUCTION

Bovine viral diarrhoea (BVD) is recognised as one of the most important endemic diseases in cattle, with a significant economic impact on the industry. This bovine pestivirus is distributed worldwide, with only a few European countries having eradicated the virus (1).

Bovine viral diarrhoea virus (BVDV) infection is usually not noticed by farmers, except in the case of rare virulent outbreaks. In most cases, the clinical presentation is non-specific, limited to a few days of fever and loss of appetite. Once a herd is exposed to BVDV, reproductive problems caused by the disease can occur, including transient infertility, abortion, stillbirths, malformed calves, and persistently infected (PI) calves. Other manifestations of the disease on herd health can also be detected, e.g., a drop in milk production, clinical mastitis, and an increase in respiratory and enteric diseases (2).

BVD control programs around the world have been developed for the prevention and control of the disease through vaccination. Vaccination against BVDV is an effective tool for disease control, as it reduces the risk of reinfection in the herd (3) as well as the likelihood of foetal infection (4). However, the use of BVDV vaccines raises concerns about interference, i.e., the induced antibody response impairing the interpretation of serological surveillance in the herd (5). The use of laboratory diagnostic assays to detect BVDV-specific antibodies in individual samples of serum and milk, as well as in bulk tank milk, is often part of BVD control programs, in terms of both serological surveillance and estimating BVDV prevalence in the herd. Detection of the virus (antigen ELISA or real-time reverse transcriptase-PCR) is used to identify PI animals in order to cull them and reduce the risk of infection within the herd (6).

The structural proteins of BVDV are the nucleocapsid and the three envelope glycoproteins E^rns^, E1 and E2 (7). The structural envelope glycoprotein E2 is the major immunogenic determinant of the BVDV virion (8). Neutralising antibodies (SN) induced in infected animals are mainly directed against E2 (9). Moreover, E2-specific monoclonal antibodies can neutralise both BVDV-1 (pestivirus A) and BVDV-2 (pestivirus B) (10).

The non-structural catalytic serine protease (NS3 or p80) is a highly immunogenic protein from BVDV and is the basis of several commercially available ELISAs. Some authors (11) showed that, after the primary courses of inactivated BVDV vaccines, antibodies against p80 protein were not induced. However, other studies (3, 12, 13, 14) showed inconclusive results regarding the reliability of different commercial p80 ELISAs to distinguish vaccinated from infected animals.

A novel BVDV subunit vaccine has been developed (DIVENCE®; 15), comprising different bovine respiratory and reproductive antigens, including BVDV-1 E2 recombinant glycoprotein, and BVDV-2 E2 recombinant glycoprotein.

Recently, the efficacy of DIVENCE® has been demonstrated against experimental challenges of BVDV-1 and BVDV-2 in pregnant animals (16). The immune response of those animals was assessed, with high titres of total antibodies (ELISA) and neutralising antibodies against BVDV-1 and BVDV-2 being described on the day of challenge. Challenge trials cannot be conducted for all potential infection situations, such as days after vaccination or days of gestation. Furthermore, animal rights concerns prevent their widespread use. For this reason, antibody analysis is a useful method to assess the immune responses induced by the vaccines (8).

The design of this E2-recombinant protein-based BVDV vaccine allows infected animals to be differentiated from vaccinated animals (DIVA, or marker vaccine). DIVA vaccines have been used in the veterinary field for decades. They carry at least one antigenic protein less than the wildtype virus; the diagnostic test thus measures the antibodies against the absent protein(s) to identify infected animals (17).

The purpose of the study was to determine the immune response against BVDV types 1 and 2 (BVDV-1 and BVDV-2) induced by DIVENCE® as measured by different assays on serum and milk samples after vaccination, as well as to demonstrate the capacity of the vaccine to differentiate between vaccinated and infected animals.

## MATERIALS AND METHODS

### Trial 1

#### Animals and housing

Twenty young Friesian-Holstein calves (55-104 days of age) were selected from commercial farms in Catalonia (Northeast Spain). These farms had no known recent history of BVDV and had received no previous vaccination against this disease. The calves were serologically screened for BVDV antibodies (BVDV Total Antibody Test, IDEXX) and were confirmed to be free of persistent BVDV infection by real-time reverse transcription polymerase chain reaction (RT-qPCR) before being included in the study. The group size was selected in order to achieve significant differences between groups and to ensure monitoring the immune response for almost two years. This group size used was determined to contain at least eight animals during the entire study. Twenty calves were randomly divided into two study groups of ten animals (vaccinated and control group) and individually identified. The entire study was conducted on a commercial farm. All calf-handling practices were approved by the local authorities (Generalitat de Catalunya) and followed the recommendations of Directive 2010/63/EU of the European Parliament and of the Council on the protection of animals used for scientific purposes as well as the HIPRA Animal Experimentation Committee. All animals used in this study were handled in strict accordance with good clinical practice, and all efforts were made to minimise suffering. This phase was conducted over a 20-month period.

#### Vaccination and study design

All animals were given a 2 mL intramuscular dose of DIVENCE® or phosphate-buffered saline (PBS) according to the group. The vaccine was reconstituted and diluted according to the manufacturer’s instructions.

The administration regimen was conducted following the manufacturer’s instructions, i.e., two doses three weeks apart (D0 and D21 of the trial) for the basic vaccination scheme. Six months later, a booster was administered (D204); a further year later, subsequent re-vaccination (D570) was performed. During the entire study, from the first vaccination dose until 21 days after the fourth dose, all animals were monitored to assess safety parameters, such as depression and systemic reactions, as well as the immune response. Blood samples were collected from all animals to assess total antibodies against BVDV (ELISA) and neutralising antibodies against BVDV-1 and BVDV-2. These blood samples were collected prior to vaccination (D0; first dose V1) and on Days 21 (D21; second dose V2), 42, 154, 204 (D204; third dose V3), 225, 280, 395, 497, 570 (D570; fourth dose V4) and 591 of the study. Whole blood samples were also collected prior to vaccination (D0) to obtain buffy coats in order to assess the absence of BVDV virus before the study began.

#### Laboratory analysis

All serum samples were analysed using a commercial ELISA (BVDV Total Antibody Test, IDEXX; ELISA A). This ELISA is based on the use of antigens derived from whole BVDV. Serum samples were also analysed using a neutralisation test in MDBK cells in 96-well plates to quantitate SN antibodies against BVDV-1 and BVDV-2. Singer (BVDV-1) and VV-670 (BVDV-2) cytopathic virus strains were used to perform the neutralisation test. A constant viral titre (10^2^ CCID_50_) was incubated with two-fold dilutions of sera. Culture plates were incubated for seven days and visually assessed for virus-induced cytopathic effects.

### Trial 2

#### Animals and study design

Three commercial dairy farms from Spain were selected (farms TON, SON, SUB), which were all seronegative for BVDV at the start of the study. For each herd, a sample of bulk tank milk before vaccination was also collected and analysed by PCR and ELISA (CIVTEST Bovis BVD/BD p80; HIPRA) by an external laboratory, ALLIC (Associació Interprofessional Lletera de Catalunya); all farms had a negative result.

The study was designed as a multicenter, randomised, placebo-controlled, double-blind field trial comparing the DIVENCE® vaccine to a placebo (phosphate-buffered saline; PBS). On each farm, animals were randomly allocated into two groups on D0, the DIVENCE® or control group, stratified by animal category (heifer or cow). In the heifer category, animals from 10 months of age until calving were included, and in the cow category, animals between first and seventh calving were included. Around half of the animals were vaccinated with DIVENCE® and the other half received PBS (SON: 260 control and 258 vaccinated animals; SUB: 73 control and 75 vaccinated animals; TON: 191 control and 198 vaccinated animals), using the same volume and the same days described for Trial 1.

Blood samples were collected from 32 random animals on each farm (16 vaccinated and 16 control animals) immediately prior to vaccination (D0) and on Days 21, 42, 204, 225, 385, 570 and 591 of the study. Additionally, bulk milk samples from each farm were obtained prior to vaccination and approximately 12, 15, 18, 21 and 24 months after the first dose of vaccine (D0). Additionally, on farm TON, individual milk samples were collected from 58 milking cows (27 vaccinated and 31 control) on D591 (21 days after the fourth vaccination).

This trial was approved by the Spanish National Agency for Medicines and Medical Devices (AEMPS; License no. 21/010). The trial was conducted in compliance with the Good Clinical Practice Guidance Document (18).

#### Laboratory analysis

Three different commercial ELISA tests were used in this trial. The BVDV Total Antibody Test (IDEXX) is based on use of antigens derived from whole BVDV (ELISA A, total antibodies). The other two ELISA tests were specifically based on detection of the p80 antigen (BVDV NS3) of the virus, CIVTEST Bovis BVD/BD p80 (HIPRA; ELISA B) and BVDV p80 antibody test (IDEXX; ELISA C). All these ELISAs were conducted in accordance with the manufacturer’s instructions.

All serum samples were analysed with ELISA A and B. Serum samples from one farm (SON) were also analysed with ELISA C. Bulk tank milk samples were analysed using these three different commercial ELISAs, A, B and C. Individual milk samples were analysed using ELISA A and B.

Additionally, serum samples from five vaccinated animals from one farm (TON) collected on D591 of the study (21 days after the fourth dose; V4) were analysed to determine neutralising antibodies (SN). For this purpose, BVDV serum neutralising antibody detection (SN) in MDBK cells in 96-well plates was used to quantitate SN antibodies against BVDV-1 and BVDV-2. Several virus strains were used for this: Singer (BVDV-1a), SKOL 1146 (BVDV-1a), Phillips P3 (BVDV-1b), RVB1297 (BVDV-1d) and SDARS 91W, Iguazú, Ghent, VV-670 (all BVDV-2). A constant virus titre (10^2^ CCID_50_) was incubated with two-fold dilutions of sera. Culture plates were incubated for seven days and visually assessed for a virus-induced cytopathic effect in the case of cytopathic strains. For non-cytopathic strains, culture plates were incubated for four days, and an immunoperoxidase monolayer assay (IPMA) was performed.

### Statistical analysis

Statistical analyses and plots were generated using R (version 4.0.5) and Microsoft® Excel 2010 (Microsoft corp.). For the analysis of longitudinal data, mixed effects models of repeated measures (MMRM) were used. In these models, the group factor, the day factor and the group-by-day interaction were included as fixed effects. When possible, an unstructured variance-covariance matrix was employed to model within-subject errors. If convergence issues were encountered, a first-order autoregressive variance-covariance matrix was modelled instead. Models for neutralising antibody data were fitted for the vaccinated groups only from Day 21, as the other group and time conditions had all observations equal to zero. REML was used for model estimation, and multiple comparisons were adjusted using the multivariate-t distribution correction in the emmeans package of R. Total antibodies and anti-p80 antibodies were compared using a Mann-Whitney U test between groups on individual milk samples and using an ANOVA test on bulk tank milk samples. A significance level of *p*<0.05 was used for all variables evaluated in this trial.

## RESULTS

### Trial 1

Mean serum ELISA values for total antibody detection and neutralising antibodies against BVDV-1 (Singer) and BVDV-2 (VV670) for vaccinated and control animals are shown in Figure 1 (A, B, C, respectively). Vaccinated animals showed a significantly (*p*<0.05) higher titre of total and neutralising antibodies against BVDV-1 and BVDV-2 from Day 42 post-vaccination until the end of the study compared to control group. After the primary vaccination (2 doses of DIVENCE®), the antibody titre as detected by ELISA and neutralising antibodies against BVDV-1 and 2 in the vaccinated animals remained similar from D42 until the administration of the third dose (D204). After the third dose, a clear boost in antibody response was then induced, in terms of both total and neutralising antibodies against BVDV-1 and BVDV-2. These high titres remained stable during the entire year after the third dose, from D225 to D570. All vaccinated animals had neutralising antibodies against BVDV-1 from D42 until the end of the study. In terms of neutralising antibodies against BVDV-2, all animals had antibodies during the entire year after the third dose.

**Figure 1.**
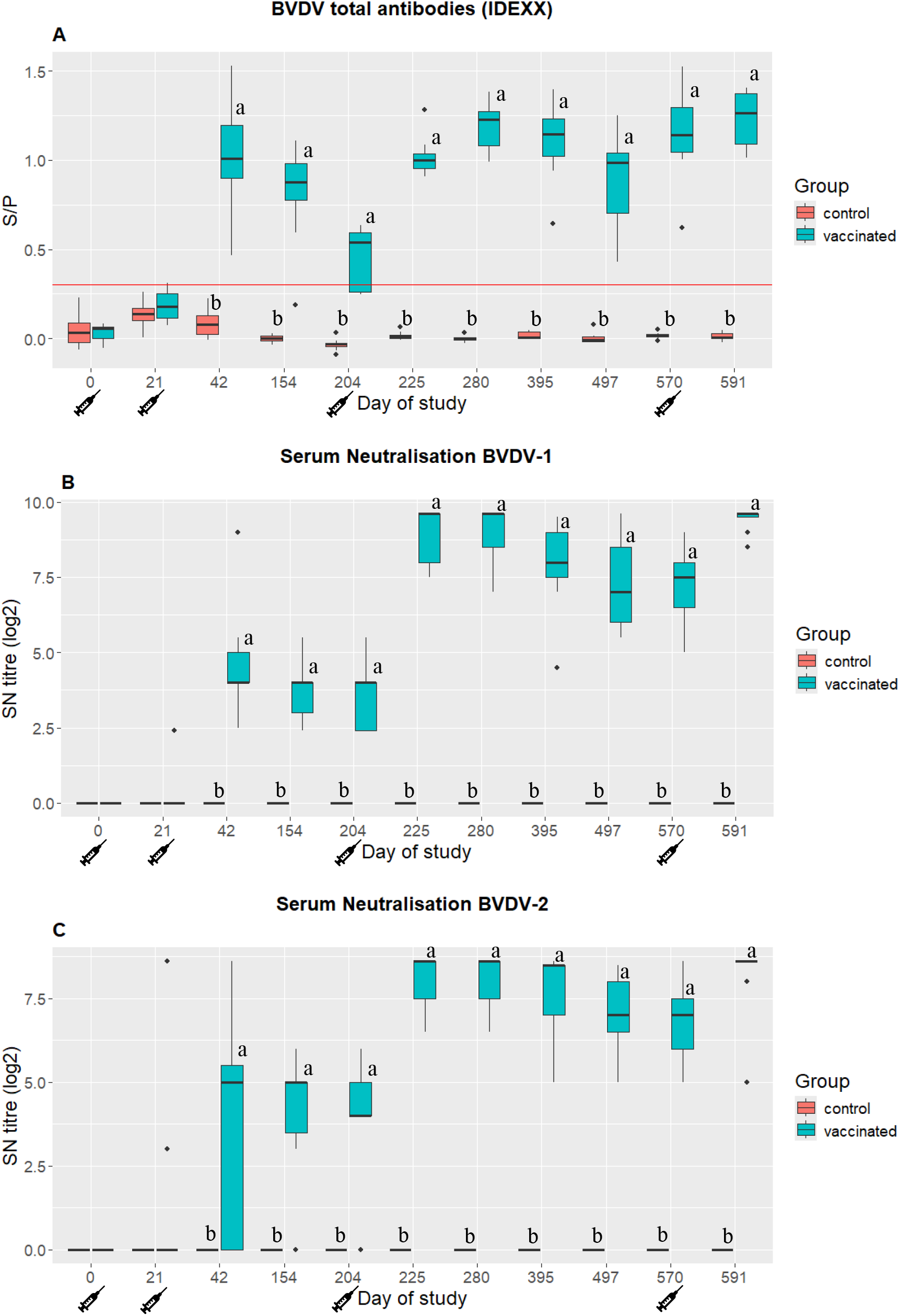
ELISA antibody titres against BVDV (A; BVDV Total Antibody Test, IDEXX), and neutralising antibodies against BVDV-1 (B; using Singer strain) and BVDV-2 (C; using VV-670 strain) from first vaccination (D0 of study) to 21 days after the one-year booster vaccination (D594 of study). ^a, b^ indicate statistically significant differences (*p*<0.05) between antibody titres in the control and vaccinated groups.

### Trial 2

#### Serology

Mean serum ELISA values for total antibody detection and anti-p80 antibody detection for vaccinated and control animals are shown in Figure 2. Vaccinated animals showed a significant (*p*<0.05) increase in total antibodies at Day 21 post-vaccination. Peak mean total antibodies were reached on D225 of the study, 21 days after the third dose of DIVENCE®. Antibody levels remained high over the entire year after the third dose, then decreased slightly prior to the fourth dose of the vaccine (D570). However, in terms of the proportion of animals with antibodies, the vast majority were positive during this period from the third dose to the fourth dose (Figure 2). Conversely, only three out of 48 control animals on one-off days had positive results for total antibodies as detected by ELISA during the entire study. Of these, none had neutralising antibodies against BVDV-1 or BVDV-2. Consequently, vaccinated animals had a significantly (*p*<0.05) higher proportion of seropositive animals compared to the control group from D21 to the end of the study (D591).

**Figure 2.**
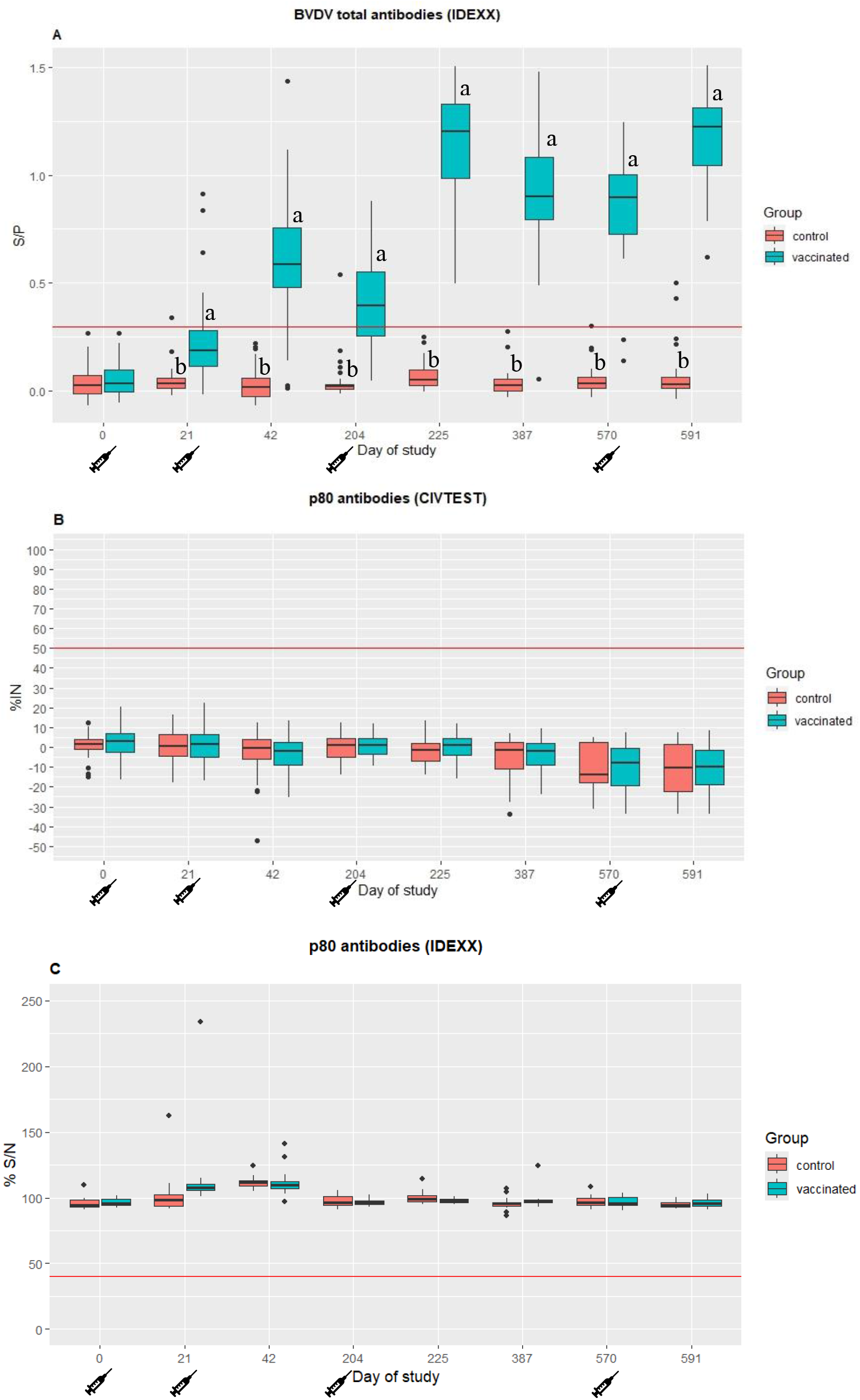
ELISA results from serum samples tested using three different kits. (A) Sample/positive control index (S/P) measured using a total antibody ELISA (BVDV Total Antibody Test, IDEXX), (B) Inhibition percentage (% IN) value measured using a p80 antibody ELISA (CIVTEST BOVIS BVD/BD p80, and (C) Signal/noise percentage (% S/N) for p80 antibody (IDEXX BVDV p80 antibody). ^a,b^ indicate statistically significant differences between groups (*p*<0.05).

In terms of anti-p80 antibodies (ELISA B and C), all animals in the vaccinated and control groups remained negative during the entire study; consequently, no differences between groups were observed.

Neutralising antibodies against several strains (BVDV-1: Singer (BVDV-1a), SKOL 1146 (BVDV-1a), Phillips P3 (BVDV-1b), RVB1297 (BVDV-1d); BVDV-2: SDARS 91W, Iguazú, Ghent, VV-670) were analysed from serum samples collected from five vaccinated animals on Day 591 of the study (Figure 3). All blood samples had neutralising antibodies against all strains listed. Mean neutralising antibody titres in these vaccinated animals were log_2_ 7.7 (range log_2_ 4-11.1) against BVDV-1 strains and log_2_ 8.0 (range log_2_ 4.5-10.1) against BVDV-2 strains.

**Figure 3.**
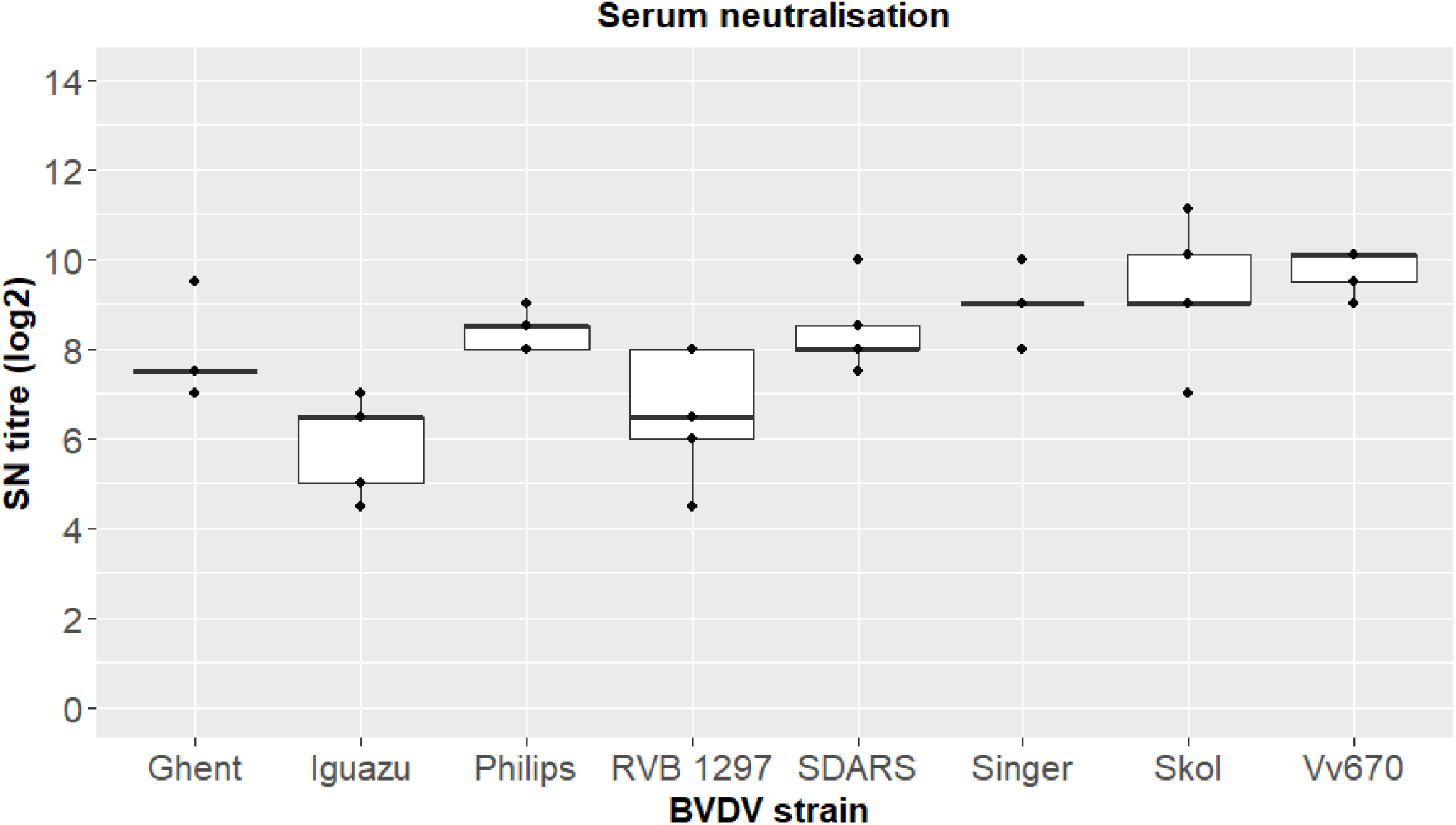
Neutralising antibodies against several strains of BVDV-1 and BVDV-2 (mean ± SD) measured by a serum neutralisation test at Day 591 of study.

#### Milk samples

Before vaccination, all bulk tank milk (BTM) samples were negative by ELISA (CIVTEST Bovis BVD/BD p80; HIPRA) conducted at the external laboratory ALLIC. After the first vaccination, total and anti-p80 antibodies were determined in BTM samples with ELISA A, B and C. No statistical differences were observed between sampling points after vaccination (12, 15, 18 21 and 24 months after the first vaccination; Figure 4) for total and anti-p80 antibodies.

**Figure 4.**
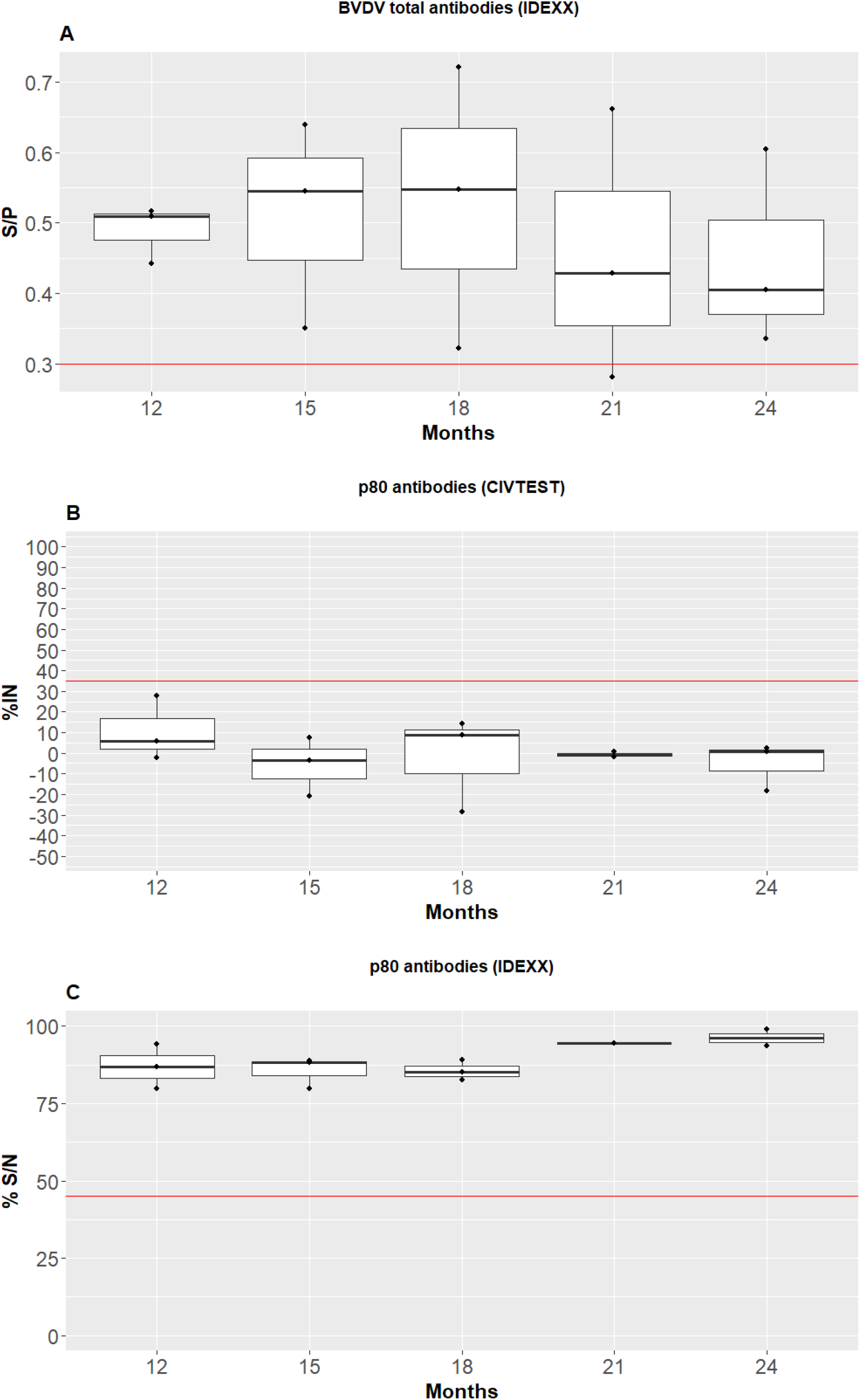
ELISA results from bulk tank milk samples tested using three different kits. (A) Sample/positive control index (S/P) measured using a total antibody ELISA (BVDV Total Antibody Test, IDEXX), (B) Inhibition percentage (% IN) value measured using a p80 antibody ELISA (CIVTEST BOVIS BVD/BD p80, and (C) Signal/noise percentage (% S/N) for p80 antibody (IDEXX BVDV p80 antibody).

In terms of total antibodies against BVDV (ELISA A), the vast majority of samples remained positive for 24 months after vaccination. Conversely, all BTM samples were negative for anti-p80 antibodies (ELISA B and C) from 12 to 24 months after vaccination.

Individual milk samples collected 21 days after the fourth vaccination were also analysed using ELISA A and B. In terms of total antibodies against BVDV, 26 out of the 27 vaccinated animals were seropositive, while all control animals were seronegative. Consequently, vaccinated animals had significantly (*p*<0.05) higher total antibody titres against BVDV compared to control animals (Figure 5). At the same time, anti-p80 antibody levels remained negative in both groups, and no statistically significant differences were observed between control and vaccinated animals.

**Figure 5.**
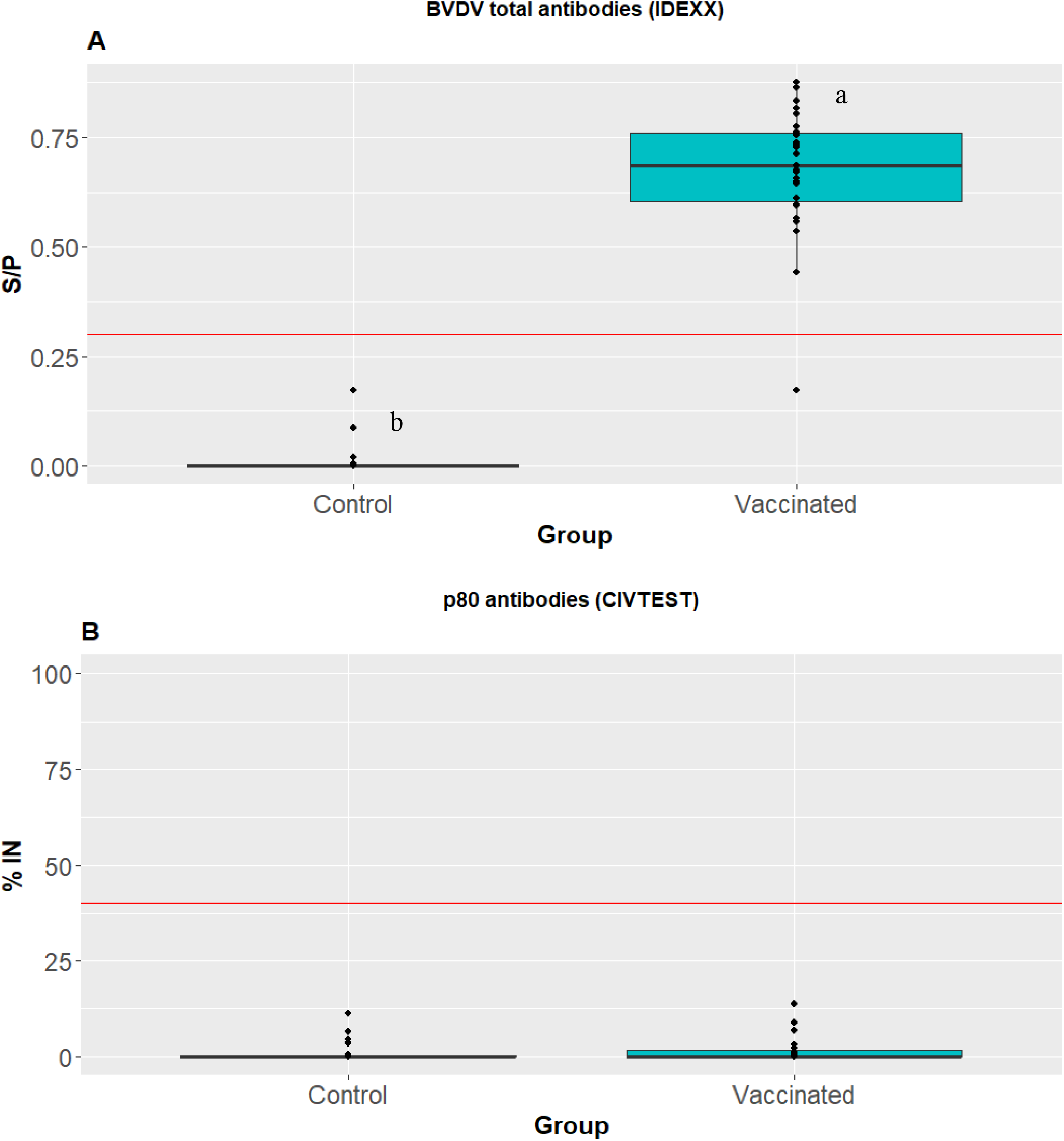
ELISA results from individual milk samples from vaccinated and control animals at Day 591 of the study, tested using two different kits. (A) Sample/positive control index (S/P) measured using a total antibody ELISA (BVDV Total Antibody Test, IDEXX), (B) Inhibition percentage (% IN) value measured using a p80 antibody ELISA (CIVTEST BOVIS BVD/BD p80). ^a,b^ indicate statistically significant differences between groups (*p*<0.05).

## DISCUSSION

The aim of this article was to investigate the immune response against BVDV types 1 and 2 (BVDV-1 and BVDV-2) induced by DIVENCE® as well as its suitability as a DIVA vaccine. To do so, two trials were conducted.

In the first trial, the vaccine resulted in a clear increase in antibodies, both in terms of ELISA (total antibodies) and neutralising antibodies. The immune response remained stable for 12 months after the third dose, with no statistical differences found between sampling points (from D225 to D570 of the study). Given the characteristics of dairy farms, with animals at different production stages across the year, it must be ensured that all animals are sustainably and effectively protected throughout the production cycle. If infection with non-cytopathic BVDV strains occurs during the first 125 days of gestation, the foetus can become persistently infected (PI) with the virus (18). Therefore, a high and sustained response over time is fundamental as a tool to fight against transplacental BVDV infection.

The efficacy of DIVENCE® against transplacental infection of BVDV-1 and BVDV-2 in pregnant animals has been demonstrated in a previous study (16). In that study, DIVENCE® was also found to induce both a marked humoral and cellular immune response, increasing the level of antibodies and interferon gamma. The results obtained in that study on neutralising antibody titres in vaccinated animals on the day of both challenges were statistically similar to the levels obtained in the present study at all sampling points from D225 to D570. These results show that the immune response induced by DIVENCE® after the re-vaccination scheme confers high and stable levels of neutralising antibodies against BVDV-1 and BVDV-2 able to reduce transplacental infection for one year.

In Trial 2, the humoral response of DIVENCE® was evaluated on serum and milk samples (BTM and individual) with different ELISA kits, one based on the use of antigens derived from whole BVDV (ELISA A) and two specifically based on anti-p80 antibodies (BVDV NS3, ELISA B & C). The vaccine induced a strong increase in total antibody levels from Day 21 of the study but left anti-p80 antibody titres negative on serum and milk samples, even after subsequent re-vaccinations. In terms of milk samples, only one vaccinated animal had no anti-BVDV antibodies in the milk, equivalent to 3.7% of the vaccinated animals. This may be due to different causes, such as the sensitivity of the IDEXX Total Antibody Test (96.3%), the individual response which may be affected by the state of the animal the day it received the fourth dose, or influences such as high milk production, which could dilute the antibody titre in the milk. In any case, it should be noted that, while vaccination efficacy may depend on the individual (animal), it is effective at the herd level (8, 19, 20). The use of safer vaccines such as DIVENCE® allows mass vaccination strategies, which have been demonstrated to increase the success of vaccination programs by enhancing herd immunity (19, 20).

The structural envelope glycoprotein, E2, is the major immunogenic determinant of the BVDV virion (8). Neutralising antibodies (SN) induced in infected animals are mainly directed against E2 (9). Moreover, monoclonal antibodies specific to E2 have demonstrated a virus-neutralising ability against both BVDV-1 and BVDV-2 (10). These findings support the design of a BVDV vaccine based on E2 to distinguish infected from vaccinated animals. Accordingly, the E2 protein has been suggested as a promising approach for a subunit vaccine (21). Different studies have been published using this approach towards future vaccine candidates (22, 23, 24, 25, 26, 27, 28).

At the same time, since non-structural proteins are mainly produced during viral replication, cattle develop antibodies to these antigens following natural infection and/or vaccination with MLV vaccines. The interference of MLV vaccines in diagnostics goes beyond antibody detection. The strains contained in the MLV vaccines have been found to be detectable by PCR in ear notch samples of newborn calves (29), requiring additional analysis to differentiate infected from vaccinated calves. Consequently, inactivated BVDV vaccines, where the virus cannot replicate, have been suggested to have an advantage over modified-live-virus vaccines in that they may not induce detectable antibodies against the BVDV p80 protein. However, the use of the antibodies against p80 to monitor BVDV circulation in herds after vaccination with inactivated vaccines has failed; false positive results are often obtained, and the use of inactivated vaccines has been shown to interfere with ELISA monitoring in blood and milk samples (3, 12, 13, 14).

The existence of this subunit vaccine, which induces antibodies against the E2 protein only, brings the characteristic of a real DIVA vaccine and solves these limitations.

The DIVENCE® vaccine induced neutralising antibodies against different BVDV strains. Four strains from BVDV-1 and four strains from BVDV-2 from countries in Europe and the Americas were selected to represent the antigenic diversity of this virus worldwide. Serum from all vaccinated animals was found to neutralise all strains tested. It was thus confirmed that animals vaccinated with DIVENCE® produce neutralising antibodies against a large range of BVDV strains, including genotypes 1 and 2, as well as different subgenotypes.

A comparison of the neutralising antibodies titres obtained in this study with those induced by other vaccines is out of the scope of this trial. As tests are not standardised between laboratories, a reliable comparison of this kind would be difficult. The technique depends on the cell system and the strain of virus used (30). However, a comparison between the neutralising antibody titres obtained in this trial and those obtained in previous trials using the same standardised test is possible. The neutralising antibody levels achieved against all strains tested are higher than those obtained against the Iguazú strain, which is the challenge strain used in a previous study (16). Overall, the results presented here suggest that DIVENCE® induces a protective response against the most prevalent subgenotypes of BVDV-1 and BVDV-2.

## CONCLUSION

DIVENCE® combines the immunogenicity of live-attenuated vaccines with the safety of inactivated vaccines. This vaccine provides a strong and sustained immune response against the most prevalent subgenotypes of BVD-1 and 2 (cross-protection). Compared to ordinary BVDV vaccines, DIVENCE® also allows infected animals and vaccinated animals to be differentiated (DIVA, or marker vaccine), making it a new tool to control and monitor the BVDV status of farms.

## Acknowledgments

The authors would like to thank Manuel Cañete for the statistical help analysing the data of this paper, as well as all personnel at HIPRA Scientific, particularly Sara Baila, Laura Feixas and Ricard Martín, for the help performing the study.

